# Identification of Genomic Loci Associated with Cellular Rhythms in Diversity Outbred Mice

**DOI:** 10.64898/2026.05.28.728467

**Authors:** Chen Fu, Sam-Moon Kim, Vivek M. Philip, Leona H. Gagnon, Colleen A. McClung, Elissa J. Chesler, Ryan W. Logan

## Abstract

Inter-individual variation in human molecular and behavioral circadian rhythms motivates genetic dissection in model systems with human-like diversity. We quantified cellular clock phenotypes from primary skin fibroblasts of several hundred Diversity Outbred (DO) mice-each carrying a unique mosaic of eight founder genomes-by longitudinal bioluminescence recordings of a Bmal1-luciferase reporter (LumiCycle). Canonical rhythm parameters (period, phase, amplitude, damping) were extracted and exhibited broad variability (heritability ≈13–35%), exceeding the ranges of founder strains. We performed genome-wide QTL mapping with R/qtl2 (linear mixed models with sex and experimental group covariates, kinship control, permutation-based significance, 1.5-LOD support intervals). A suggestive QTL for amplitude localized to chromosome 12 (LOD 6.9; ∼7.5–12.0 Mb), with founder effects indicating higher amplitude for C57BL/6J and lower for PWK/PhJ. Among 21 protein-coding genes in this interval, *Apob* (apolipoprotein B), a clock-regulated determinant of lipoprotein assembly, emerged as a strong candidate for amplitude control. A phase QTL mapped to chromosome 1 (support interval ∼1.36 Mb) with divergent founder effects (C57BL/6J, NOD/ShiLtJ, WSB/EiJ: delayed; PWK/PhJ, CAST/EiJ: advanced) and prioritized candidates including *Epha4* (an Eph receptor tyrosine kinase implicated in photic entrainment) and *Acsl3*. Integrative analysis in GeneWeaver connected QTL gene sets to prior loci for voluntary alcohol consumption and circadian period on proximal chromosome 12, and highlighted overlaps with GWAS signals for adolescent idiopathic scoliosis and schizophrenia, suggesting shared pathways between circadian regulation, metabolism, and neurobehavioral traits. Together, these findings define reproducible genomic loci for cellular clock phenotypes in a highly recombinant population, nominate tractable candidate genes (*Apob, Epha4/Acsl3*) for mechanistic follow-up, and illustrate how high-diversity mouse genetics bridges cellular circadian variation with complex disease biology.

## Introduction

In humans, there is substantial variation in molecular, cellular, and behavioral rhythms. A minimally invasive approach for measuring molecular rhythms in humans is the collection of skin biopsies for the culture of skin fibroblasts. Skin fibroblasts are transfected with fluorescent reporter constructs to measure molecular rhythms across days. Canonical rhythm parameters of molecular readouts can be assessed, including period, phase, amplitude, among others. Using similar approaches, certain patterns of cellular rhythms have been correlated with mood and substance use disorders. Associations between molecular rhythm phenotypes and psychiatric disorders suggest common or similar genetic contributions to circadian rhythm phenotypes and psychiatric disorders. Indeed, prior work has shown a robust relationship between circadian rhythm disruptions and vulnerability to and worsening of alcohol use disorder and other substance use disorders (Logan & McClung, 2019).

Akin to humans, broad variation of circadian rhythms is observed in several genetically diverse rodent populations. For example, circadian rhythms in fibroblasts and locomotor behavior are distributed across a wide range in the Diversity Outbred (DO) mouse population (Saul et al., 2019). The DO mouse population captures ∼45-million unique polymorphisms and allelic combinations, and several million structural variants, driving genetic diversity and expanding phenotypic heterogeneity. DO mice are bred from eight different founder strains, five inbred strains and three wild-derived strains, that provide a heterogeneous population that more closely mirrors the diversity found in human populations. This high genetic diversity is crucial for capturing the broad spectrum of phenotypic expression necessary for understanding how genetic variations contribute to complex traits, enhancing the translatability of findings to human health and disease.

The advantages of experimental use of DO mice extends to their utility in genetic mapping studies, where they offer refined resolution compared to inbred strains and other hybrid mouse panels. Each DO mouse carries a unique combination of alleles from the eight founders, resulting in a high number of genomic recombination events per mouse. Fine-scale mapping capabilities are essential for identifying a specific quantitative trait locus (QTL) associated with complex traits, such as circadian rhythms. Further, the ability to estimate allele effects from multiple founders can help refine the identification of potential candidate genes, significantly reducing the number of putative loci for complex behavioral traits.

In prior work, we demonstrated heterogeneity of molecular rhythms in cultured skin fibroblasts between inbred and wild-derived strains of mice (Kim et al., 2021). Additionally, we found far greater variability of cellular rhythms in a large sample of DO mice relative to the eight founder strains, suggesting genetic diversity leads to expansion of phenotypic heterogeneity in cellular circadian rhythms. Here, we used genome wide association analyses to identify QTL associated with key cellular rhythm phenotypes in several hundred DO mice.

## Methods

### Animals

Male and female (8-12 weeks old) mice from Diversity Outbred population and the founder strains (A/J, C57BL/6J, 129S1/SvImJ, NOD/ShiLtJ, NZO/HlLtJ, CAST/EiJ, PWK/PhJ and WSB/EiJ) were used for isolation of fibroblasts, genotyping, and genetic analyses. The complete molecular and behavioral datasets were published elsewhere (Kim et al., 2021). Mice were housed at the Jackson Laboratory under a standard 12:12-h light-dark cycle (lights on at 0600-h). All mice were fed food and water ad libitum. All animal procedures used in this study abide by the ARRIVE (Animal Research: Reporting In Vivo Experiments) guidelines and were conducted in accordance with the National Institute of Health guidelines for the care and use of laboratory animals and approved by the Institutional Animal Care and Use Committee of the University of Pittsburgh and The Jackson Laboratory.

### Genotyping of Diversity Outbred Mice

Tails were removed from each mouse at euthanasia, placed into 1.5mL Eppendorf tubes, and stored in saline at −80C until DNA extraction. Tail samples were shipped to GeneSeek (Neogen Inc.) for DNA extraction and genotyping on the GigaMUGA Illumina array platform. GigaMUGA arrays assess 143,259 genetic markers spanning 19 autosomes and the X chromosome of the mouse, with mean spacing of 18Kb. Markers were optimized for DO mice. Genotypes were imputed to a 69K grid to enable equal representation across the genome. No samples were excluded for quality control failures (missing genotypes or errors, sample duplicates, and chromosomal congruence to known sex).

### Primary Fibroblast Isolation for Cellular Rhythm Analyses

Skin biopsies from the ear (1mm diameter) were collected from mice at the Jackson Laboratory then sent to the University of Pittsburgh in Dulbecco’s Modified Eagle’s Medium (DMEM). Biopsies were digested in DMEM containing 2.5mg/ml collagenase D (Gibco) and 1.25mg/ml pronase (Millipore) for ∼90-mins. The biopsy was then plated on a 60mm tissue culture dish, overlaid with sterilized square cover glass (22mm) and containing DMEM solution (10% fetal bovine serum; 1% amphotericin B; and 0.1% gentamycin). Tissue culture dishes were maintained in an incubator at 37C°. Cultures were transfected with 1×10^7^ units of lentiviral construct—luciferase reporter fused to *Bmal1* (AddGene). Individual cultures were synchronized by bath application of 15uM forskolin (Sigma) for ∼2-h. Cells remained in DMEM supplemented with 15uM forskolin, 25mM HEPES, 292ug/ml L-glutamine, 100units/ml penicillin, 100ug/ml streptomycin, 10uM luciferin (Promega). Bioluminescence of *Bmal1-*luciferase (*Bmal1-luc*) transfected cell cultures was recorded by an automated 32-channel luminometer (LumiCycle by Actimetrics) in an incubator. Recording was conducted across 2-4 fibroblast cultures per mouse biopsy sample for ∼6-days every 70-secs at 10-mins intervals.

Primary circadian rhythm measures from luciferase reporter traces were determined using LumiCycle Analysis software, including period, amplitude, phase, and damping rate. Briefly, period was estimated as the peak-to-peak interval, while peak-to-trough was used as amplitude. The specific time when reporter activity peaked was used as phase. Damping rate was estimated as the number of days for the rhythm amplitude to decrease to ∼36% of the initial amplitude.

### QTL Mapping of Cellular Circadian Rhythm Phenotypes in Diversity Outbred Mice

Each of the major circadian rhythm phenotypes from the fibroblast rhythms were mapped using the genotypes of each DO mouse. Genome reconstruction of the DO, sample and marker quality control and mapping of QTL was completed by R/qtl2 software (v0.38), as described previously (Broman et al., 2019). R/qtl2 software is designed for QTL mapping in multi-parent populations derived from multiple founder strains. R/qtl2 performs genome scans using linear mixed models to account for population structure and impute SNP markers based on the genomes of the founder strains.

A mixed model included both sex and experimental group as additive covariates and a random effect to account for kinship. Significance thresholds were obtained by performing 1,000 permutations of the genome scans with phenotypic data shuffled among the individual mice. From the linear model, 1.5 LOD support intervals were determined for significant (<0.05) and suggestive (<0.1(probable) or <0.37(possible)) QTL peaks (Figure 1B). The filter we applied for was 7.03 for period, 6.78 for phase, 6.79 for amplitude and 8.18 for damping, corresponding to the possible (63rd percentile) thresholds in Figure 1B; the loci reported below therefore represent suggestive QTL. Regression coefficients for additive effects of alleles associated with the founder strains were estimated at each genomic locus. Plausible candidate genes were identified using functional and protein coding genes within the specified QTL intervals using R/qtl2. To further narrow candidate genes, a literature review was conducted using PubMed to identify known associations between genes and particular circadian rhythm phenotypes.

**Figure 1.**
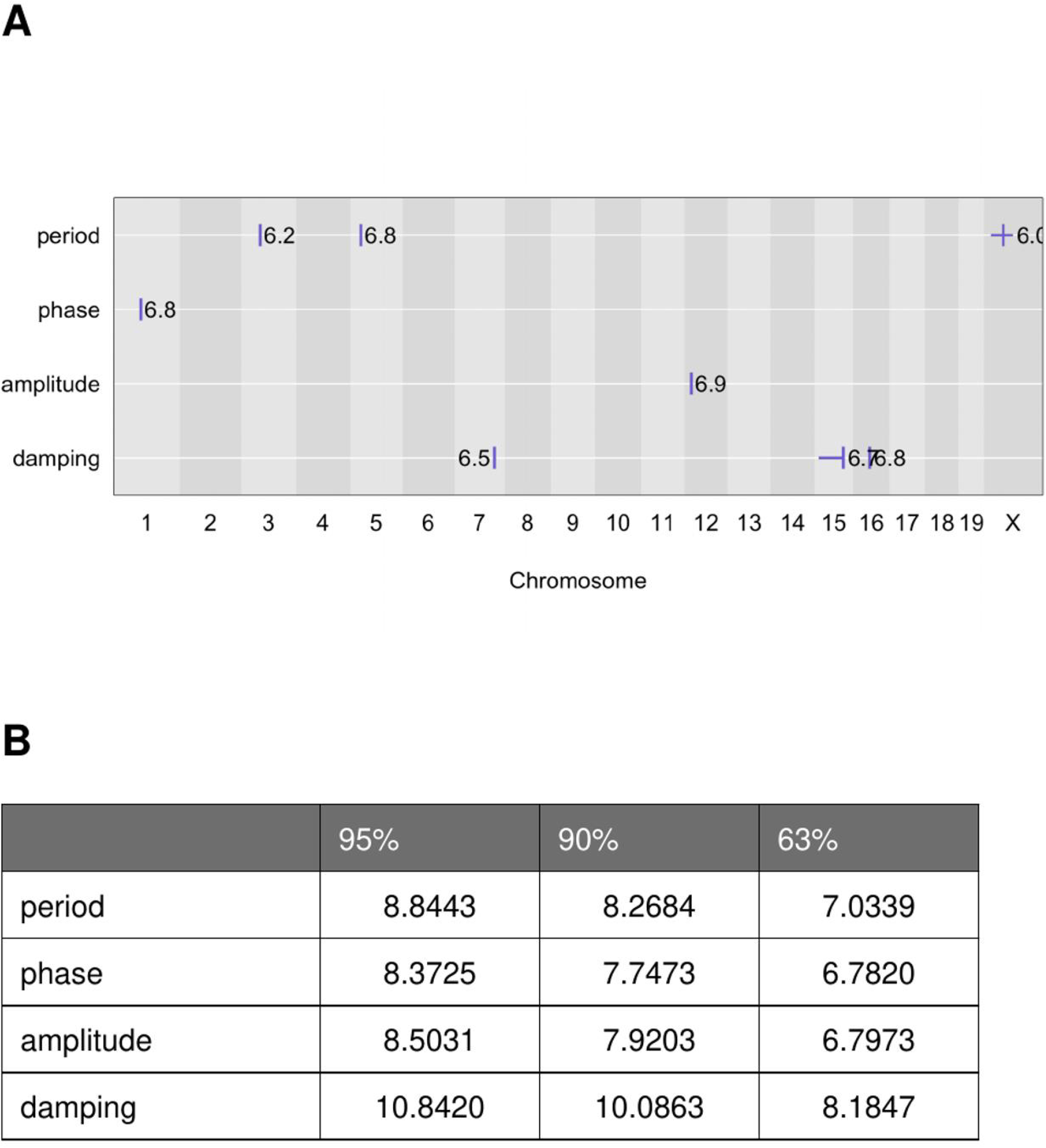
Genome-wide QTL summary and permutation-based significance thresholds for four cellular circadian phenotypes. (**A**) Locations of the QTL peaks detected for each circadian rhythm parameter — period, phase, amplitude, and damping rate — across the 19 autosomes and the X chromosome. Each tick marks the genomic position of a peak and the adjacent number gives its LOD score: period on chromosomes 3, 5, and X; phase on chromosome 1; amplitude on chromosome 12; and damping rate on chromosomes 7, 15, and 16. (**B**) Genome-wide LOD significance thresholds for each phenotype, estimated from 1,000 permutations of the genome scan with phenotypes shuffled across mice. Thresholds are reported at three stringency tiers: significant (95th percentile), probable (90th percentile), and possible (63rd percentile). The amplitude (chromosome 12) and phase (chromosome 1) peaks analyzed in subsequent figures reach the possible tier (LOD ≈ 6.80 and 6.78, respectively).

### Integrative Genomics Using GeneWeaver.Org

*GeneWeaver*’s database contains numerous gene expression datasets based on several tiers of curation confidence (Baker et al., 2012). The database was queried (July 2025) for level “Tier II” of curation for similar gene sets to the comprehensive protein coding gene set identified from each QTL. Using Jaccard Similarity, datasets with similarity scores above 0.15 were then integrated into a project that contained the combined gene set from the circadian rhythm QTL. A GeneSet graph was generated to visualize high-degree genes with high connectivity. Integration identified genes with experimental relationships between circadian rhythms and other phenotypes.

### Statistical Analysis

We used RStudio (v4.4.2) for analyses conducted via R/qtl2 with settings “qtl2”, “readxl”, “tidyverse”, and “qtl2convert”. *Metascape* (Zhou et al., 2019) was applied to find significant biological processes enriched by genes underlying each identified QTL loci.

## Results

In our prior work (Kim et al., 2021), cellular rhythm phenotypes were highly variable and extended beyond the minimum and maximum range observed in the founder strains of the DO mice. In addition, each of the cellular rhythm phenotypes had modest heritability estimates (damping rate, 13%; phase, 26%, amplitude, 26%; goodness of fit, 33%; and period, 35%). Increased heterogeneity of cellular rhythm phenotypes in the DO mice combined with identified genotypes of each DO mouse enabled us to conduct QTL mapping of cellular rhythm phenotypes.

### QTL Mapping of Circadian Amplitude

In fibroblasts from DO mice, QTL mapping identified a suggestive QTL for rhythm amplitude on chromosome 12 that surpassed the possible (63rd percentile) threshold (LOD: 6.9 vs. threshold 6.79; Position: 11.69 Mb; Figure 2A). The QTL for circadian rhythm amplitude captured an interval between 7.52 and 12.01 (4.58 Mbp). Founder allele effects showed C57BL/6J allele contributed to a higher amplitude compared to other founder alleles, while the PWK/PhJ allele was associated with lower amplitude (Figure 2B, 2C). We collected functional RNA and protein coding genes within the interval (Figure 2D). The QTL interval contained 86 genes (Table S1, S4), with 21 protein coding genes. Of these 21 genes, only *Apob* had direct links to circadian rhythms. *Apob* encodes apolipoprotein B, a key component of lipoproteins involved in lipid metabolism and transport, with known circadian regulation by core circadian genes *BMAL1* and *CLOCK* (Lee et al., 2012; Pan et al., 2010; Pan & Hussain, 2007). Another gene, *Vsnl1*, coding Visinin-like (neuronal calcium-sensor) proteins have been implicated in circadian gene expression in the pineal gland (Braunewell & Szanto, 2009); (Link, 2004), this is suggestive but not direct evidence for *Vsnl1* function in circadian behavior.

**Figure 2.**
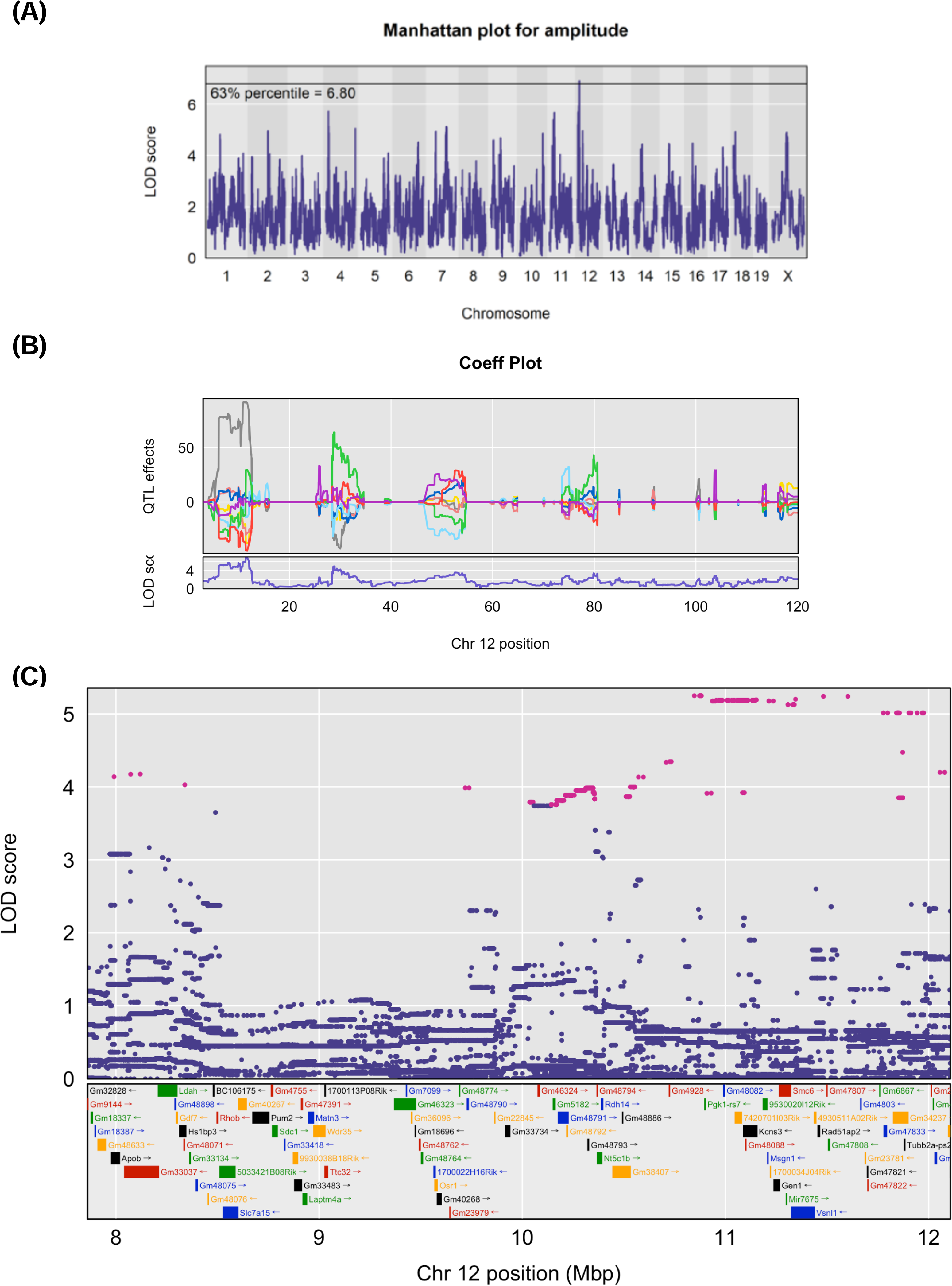
QTL for cellular rhythm amplitude on chromosome 12. (**A**) Genome-wide Manhattan plot for rhythm amplitude. The x-axis shows genomic position across chromosomes 1–19 and X; the y-axis shows the LOD score. The horizontal line marks the possible (63rd percentile) permutation threshold (LOD = 6.80). The strongest peak localizes to chromosome 12. (**B**) Founder allele effect (coefficient) plot across chromosome 12. The upper panel shows the estimated additive effects of the eight Diversity Outbred founder haplotypes on amplitude as a function of position; the lower panel shows the corresponding LOD trace. Founder color coding: A/J (black), C57BL/6J (red), 129S1/SvImJ (green), NOD/ShiLtJ (blue), NZO/HlLtJ (cyan), CAST/EiJ (magenta), PWK/PhJ (yellow), and WSB/EiJ (gray). (**C**) Fine-mapping of the chromosome 12 support interval (≈8–12 Mbp). Each point is a SNP plotted by genomic position (x-axis) and LOD score (y-axis); pink points denote SNPs within 1.5 LOD of the peak and blue points denote all other SNPs. The lower track shows the protein-coding and functional genes within the interval, with rectangles indicating gene position and length and names colored to aid legibility. *Apob*, a clock-regulated lipoprotein gene, lies within this interval.

*Geneweaver* analysis showed genes within the loci for QTL discussed above overlapped with voluntary alcohol consumption QTL 7 (Bachmanov et al., 2002) and QTL on proximal mouse chromosome 12 for inter-strain differences in circadian period (Hofstetter et al., 2007) (Figure 5B). These overlaps confirmed that our QTL for amplitude was discovered in previous circadian study. Also, it suggested functional connection between circadian behavior and alcohol consumption.

### QTL Mapping of Circadian Phase

A suggestive QTL for circadian phase mapped to chromosome 1 within a 1.36 Mb support interval (LOD: 6.75, approaching the possible [63rd percentile] threshold of 6.78; Figure 3A). Allelic effects were identified for C57BL/6J, NOD/ShiLtJ, and WSB/EiJ (delayed circadian phases), along with PWK/PhJ and CAST/EiJ (advanced circadian phases) founder strains (Figure 3B, 3C). Within the interval, there were 29 genes, including 8 protein coding genes: *Acsl3*, *Epha4*, *Farsb*, *Gm2102*, *Mogat1*, *Pax3*, *Sgpp2*, and *Utp14b* (Table S1, S5). Both *Epha4* and *Acsl3* fall within this interval, where circadian regulation of lipid metabolism is well documented (Kuang et al., 2019). *Epha4* encodes a receptor tyrosine kinase involved in modulating circadian phase and entrainment in the brain (Freyburger et al., 2016; Kiessling et al., 2018).

**Figure 3.**
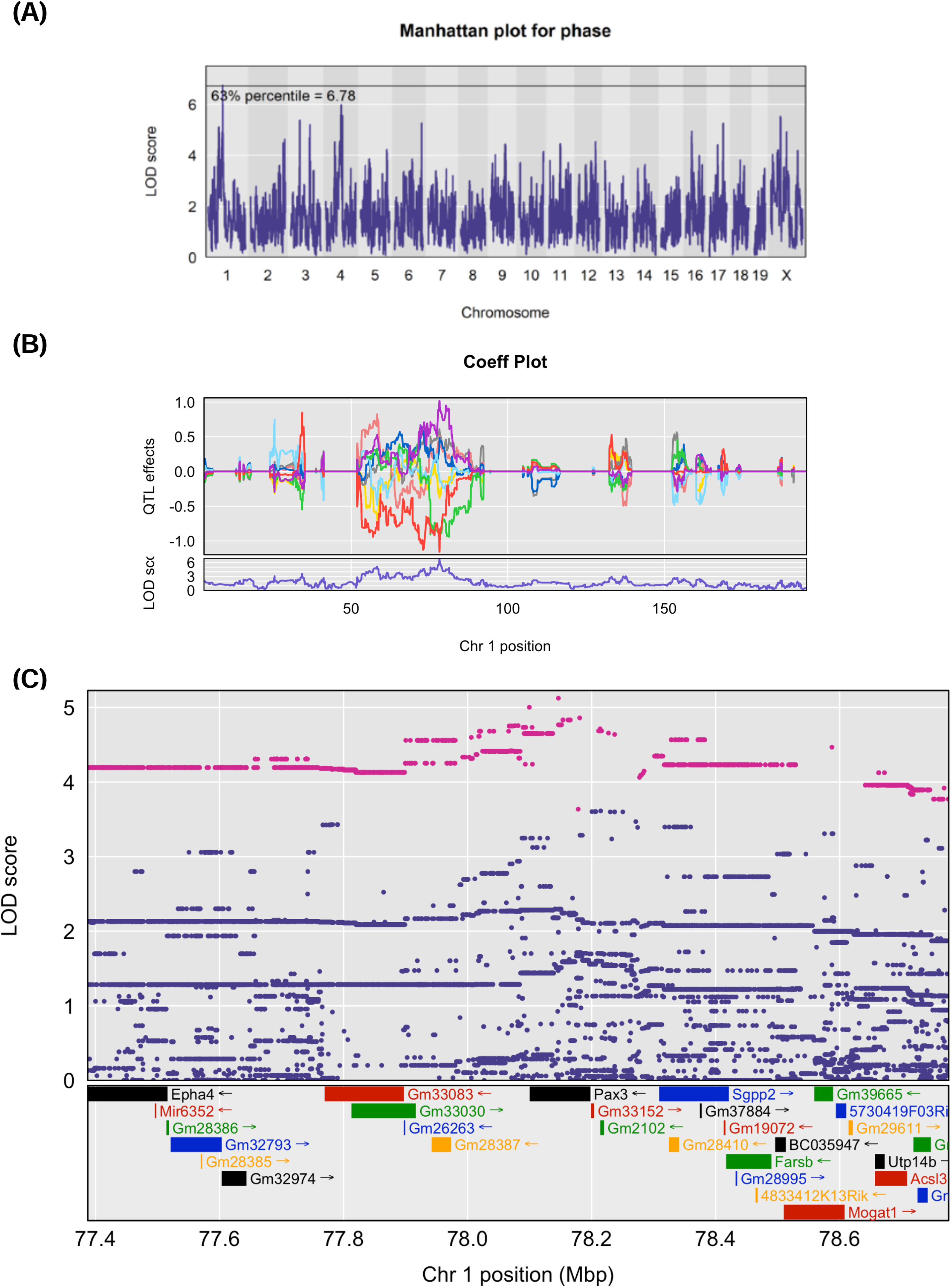
QTL for cellular rhythm phase on chromosome 1. (**A**) Genome-wide Manhattan plot for circadian phase. Axes as in Figure 2A; the horizontal line marks the possible (63rd percentile) permutation threshold (LOD = 6.78). The strongest peak localizes to chromosome 1. (**B**) Founder allele effect (coefficient) plot across chromosome 1. The upper panel shows the estimated additive effects of the eight founder haplotypes on phase as a function of position; the lower panel shows the corresponding LOD trace. Founder color coding as in Figure 2B: A/J (black), C57BL/6J (red), 129S1/SvImJ (green), NOD/ShiLtJ (blue), NZO/HlLtJ (cyan), CAST/EiJ (magenta), PWK/PhJ (yellow), and WSB/EiJ (gray). (**C**) Fine-mapping of the chromosome 1 support interval (≈77.4–78.7 Mbp). Points and color scheme as in Figure 2C: pink, SNPs within 1.5 LOD of the peak; blue, other SNPs. The lower track shows the genes within the interval — including *Epha4*, *Pax3*, *Sgpp2*, *Farsb*, *Acsl3*, *Mogat1*, and *Utp14b* — with rectangles indicating gene position and length and names colored to aid legibility.

Elucidated by *GeneWeaver*, signals under this QTL were linked to GWAS loci for adolescent idiopathic scoliosis (Kou et al., 2019) and GWAS loci for schizophrenia(Dickerson et al., 2020) (Figures 4 and 5A). This overlap suggested the link between genetic machinery underlying circadian behavior and neuropsychiatric diseases.

**Figure 4.**
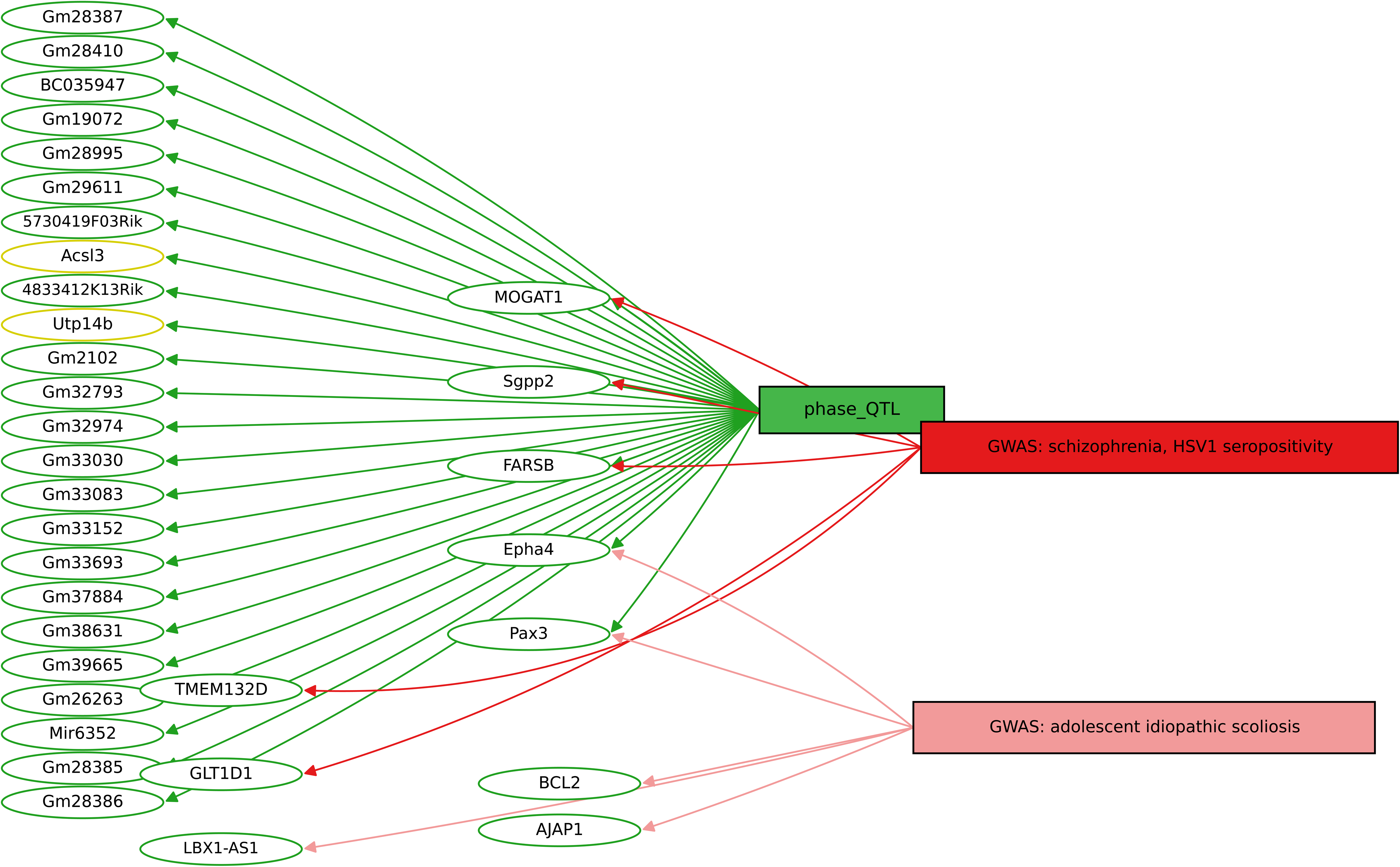
Cross-species gene–set graph for the chromosome 1 phase QTL (*GeneWeaver*). Bipartite graph relating the mouse genes under the phase QTL to overlapping human GWAS gene sets. Node coloring: green, mouse (*Mus musculus*) genes; red, human GWAS gene-set names; yellow, emphasis genes highlighted for interpretation. Genes within the phase QTL overlap gene sets for adolescent idiopathic scoliosis and for schizophrenia / herpes simplex virus type 1 (HSV-1) seropositivity, with shared candidates including the human orthologs *TMEM132D*, *GLT1D1*, *BCL2*, *AJAP1*, and *LBX1-AS1*.

### Integrative Genomics to Identify Plausible Candidate Relationships to Phenotypes

*Metascape* analysis for the 115 genes including 86 genes underlying QTL for amplitude (on chr 12) and 29 genes under QTL for phase (on chr 1) showed 4 significant GOs including alcohol metabolism (Figure. 5C, Table S6). Genes belonging to this GO are *Apob, Rdh14, Sgpp2, Mogat1. Apob* and *Rdh14* were in QTL for amplitude and *Rdh14* and *Sgpp2* were in QTL for phase. This result suggests that the circadian QTLs we found may relate to alcohol metabolism, which is consistent with the functional correlation between circadian machinery and that underlying neuropsychiatric disease, including addiction (Spanagel et al., 2005).

**Figure 5.**
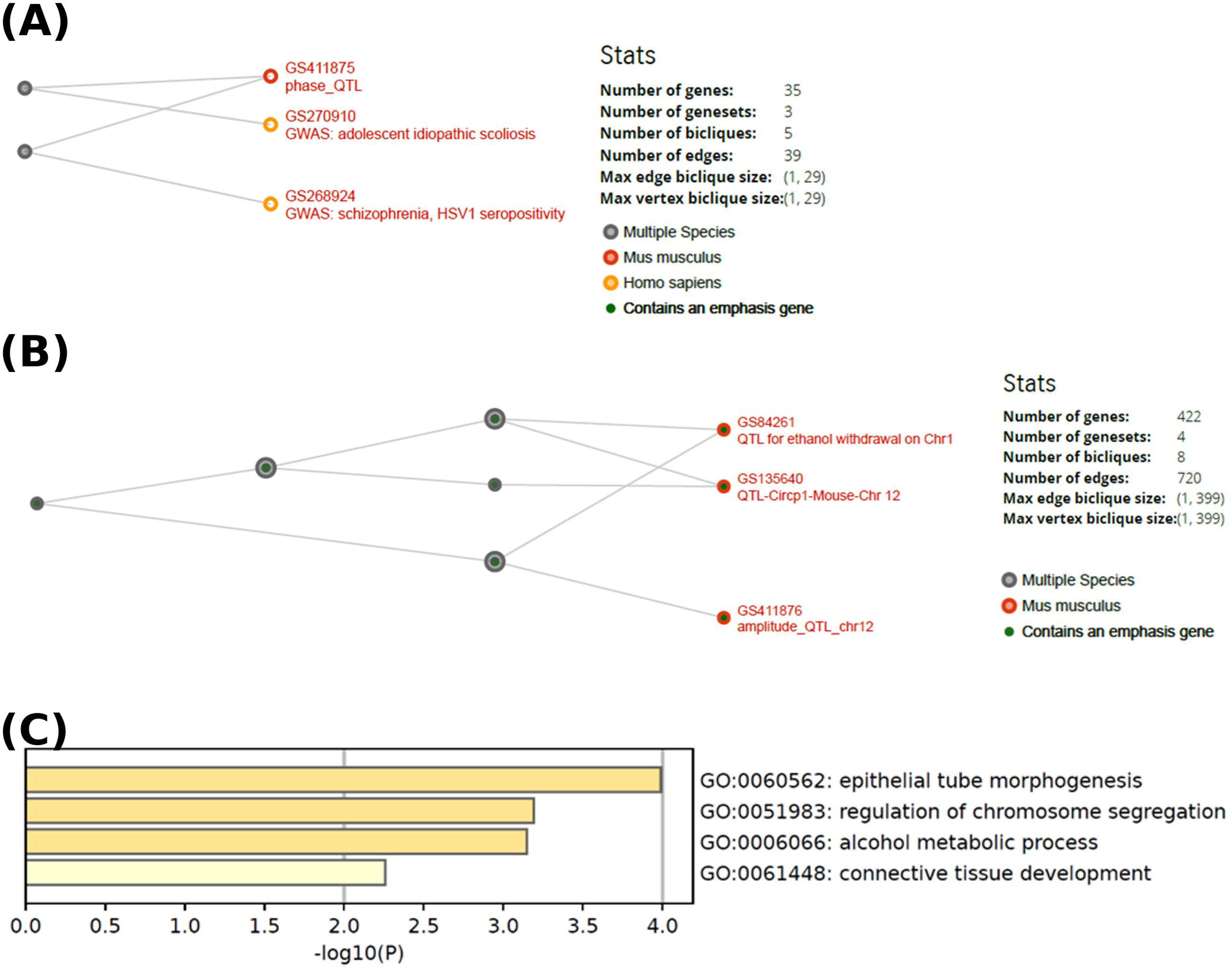
Integrative gene-set similarity and pathway enrichment for the mapped QTL. (**A**) Hierarchical similarity (HiSim) graph from *GeneWeaver* for the chromosome 1 phase QTL (GS411875, phase_QTL), showing its relationship to human GWAS gene sets for adolescent idiopathic scoliosis (GS270910) and schizophrenia / HSV-1 seropositivity (GS268924); 35 genes, 3 gene sets, 5 bicliques, 39 edges. Node coloring: gray, multiple species; red, *Mus musculus*; orange, *Homo sapiens*; a green marker indicates a node containing an emphasis gene. (**B**) HiSim graph for the chromosome 12 amplitude QTL (GS411876, amplitude_QTL_chr12), showing overlap with a QTL for ethanol withdrawal on chromosome 1 (GS84261) and a free-running circadian-period QTL on chromosome 12 (GS135640, Circp1); 422 genes, 4 gene sets, 8 bicliques, 720 edges. (**C**) Gene Ontology and pathway enrichment from *Metascape* for the 115 genes under the amplitude and phase QTL peaks. Bars show statistical enrichment as −log_10_(P) for the top terms: epithelial tube morphogenesis (GO:0060562), regulation of chromosome segregation (GO:0051983), alcohol metabolic process (GO:0006066), and connective tissue development (GO:0061448).

## Discussion

The results of our QTL mapping in DO mice revealed suggestive genetic loci associated with cellular circadian rhythm phenotypes, including amplitude, damping rate, period, and phase. These findings underscore the genetic complexity of circadian rhythms and provide insight into the molecular mechanisms underlying their regulation. The high variability in cellular rhythm phenotypes observed in DO mice, extending beyond the range of their founder strains, highlights the power of this population for dissecting complex traits (Saul et al., 2019). Our prior work indicates that measuring cellular rhythms using the canonical circadian gene *Bmal1* in skin fibroblasts is a robust and reliable approach for assessing variation in circadian rhythms in DO mice (Kim et al., 2021). Consistent with that work, cellular rhythms were highly variable between individual DO mice, and the modest heritability estimates (13–35%) suggest that while genetic factors contribute to these phenotypes, environmental or stochastic factors also play a role, consistent with the polygenic architecture of circadian rhythms. Expanding on these previous findings with updated genotypes for each individual DO mouse, we now provide high-resolution genetic mapping with combinatorial founder allelic effects for each circadian rhythm phenotype. By combining allelic effects with the sequence variation and genes within each mapped locus, we further narrowed the set of putative candidate genes within each QTL.

The suggestive QTL on chromosome 12 for circadian amplitude contained 21 protein-coding genes. The presence of *Apob* within this interval is notable given its established links to circadian biology. *Apob* encodes apolipoprotein B, a structural component of lipoproteins that is central to lipid metabolism and transport, and its expression is under circadian control by the core clock factors BMAL1 and CLOCK (Lee et al., 2012; Pan & Hussain, 2007; Pan et al., 2010). The founder allele effects, in which the C57BL/6J haplotype was associated with higher amplitude and the PWK/PhJ haplotype with lower amplitude, point to strain-specific contributions to amplitude regulation. A second gene in the interval, *Vsnl1*, which encodes a neuronal calcium-sensor protein, has been implicated in circadian gene expression in the pineal gland, although the evidence for a direct role in circadian behavior remains indirect (Braunewell & Szanto, 2009; Link, 2004). Cross-species gene-set analysis in GeneWeaver provided independent support for this locus: genes under the amplitude QTL overlapped with a previously reported QTL for free-running circadian period on proximal chromosome 12 (Hofstetter et al., 2007) and with a QTL for voluntary alcohol consumption (Bachmanov et al., 2002), indicating that this region has been associated with both circadian and alcohol-related phenotypes in independent studies. Together, these observations are consistent with reciprocal coupling between lipid metabolism and the molecular clock (Pan et al., 2010), and they nominate *Apob* as a candidate through which this coupling may influence the robustness of cellular rhythms.

The suggestive QTL for circadian phase on chromosome 1 contained eight protein-coding genes, including *Epha4* and *Acsl3*. *Epha4* encodes an Eph receptor tyrosine kinase that has been implicated in circadian clock function and in the regulation of sleep, with roles in modulating phase and entrainment in neural tissues (Freyburger et al., 2016; Kiessling et al., 2018). The founder allele effects, with the C57BL/6J, NOD/ShiLtJ, and WSB/EiJ haplotypes associated with delayed phase and the PWK/PhJ and CAST/EiJ haplotypes with advanced phase, indicate strain-specific contributions to phase regulation and are consistent with a signaling-based mechanism for clock entrainment that may involve receptor tyrosine kinase activity. Several additional genes in the interval participate in lipid metabolism, including *Acsl3* (long-chain acyl-CoA synthetase) and *Mogat1* (monoacylglycerol O-acyltransferase), echoing the lipid-metabolism theme observed at the amplitude locus. Integrative analysis in GeneWeaver linked the phase QTL to human GWAS gene sets for adolescent idiopathic scoliosis (Kou et al., 2019) and for schizophrenia and HSV-1 seropositivity (Dickerson et al., 2020), with shared candidates including the orthologs of *TMEM132D*, *GLT1D1*, *BCL2*, *AJAP1*, and *LBX1-AS1*. These overlaps suggest that the genetic determinants of cellular circadian phase intersect with pathways relevant to both skeletal development and neuropsychiatric disease.

Pathway enrichment of the combined set of 115 genes under the amplitude and phase QTL peaks (Metascape) recovered terms for epithelial tube morphogenesis, regulation of chromosome segregation, alcohol metabolic process, and connective tissue development (Zhou et al., 2019). The enrichment for alcohol- and lipid-related metabolic processes is consistent with the recurrent appearance of lipid-metabolism genes across both loci and with the overlap of the amplitude QTL with alcohol-related QTL, reinforcing a connection between circadian regulation and metabolic and substance-use phenotypes that has been described at the level of individual clock genes (Spanagel et al., 2005). The enrichment for connective tissue development and epithelial morphogenesis parallels the overlap of the phase QTL with the scoliosis GWAS, while the link to neuropsychiatric gene sets is in keeping with the broad role of circadian disruption across brain disorders (Logan & McClung, 2019). Although these cross-species and pathway associations are correlative, they are recovered consistently across independent analytic approaches and help prioritize the loci and candidate genes for mechanistic follow-up (Baker et al., 2012).

Several limitations warrant consideration. The loci reported here reached the possible (63rd percentile) permutation threshold rather than genome-wide significance and should therefore be regarded as suggestive and interpreted with caution pending replication in independent cohorts. The modest heritability estimates further indicate that non-genetic factors, including the cellular environment and stochastic processes, contribute to these phenotypes. As with any QTL study, not all genes within a support interval are causal for the phenotype, and finer mapping or functional validation is required to distinguish causal genes from linked passengers. The candidate genes nominated here were prioritized using founder allele effects, sequence variation, and prior literature, and their roles in cellular rhythm regulation remain to be tested directly. Future studies could use CRISPR-based gene editing to validate candidate genes such as *Apob*, *Vsnl1*, *Epha4*, and *Pax3*, and could integrate multi-omic data — for example transcriptomic and proteomic profiling of fibroblasts across the founder haplotypes — to define the mechanisms by which these genes shape circadian phenotypes.

In conclusion, our QTL mapping in DO mice provides a framework for understanding the genetic basis of cellular circadian rhythms. The identification of candidate genes and enriched pathways underscores the complexity of circadian regulation and its intersection with lipid metabolism, substance-use, and neuropsychiatric phenotypes. These findings provide a basis for future studies to dissect the molecular mechanisms underlying cellular circadian phenotypes and their implications for health and disease.

## Ethics Approval

All animal procedures abided by the ARRIVE guidelines and were conducted in accordance with the NIH Guide for the Care and Use of Laboratory Animals and were approved by the Institutional Animal Care and Use Committees of the University of Pittsburgh and The Jackson Laboratory.

## Author Contributions

CF and S-MK performed experiments and analyzed data; VMP, LHG, EJC, and CAM contributed to study design and data acquisition; RWL conceived and supervised the study; CF and RWL drafted the manuscript. All authors reviewed and approved the final manuscript. Confirm initials and roles.

## Data Availability

The complete molecular and behavioral phenotype datasets were previously published (Kim et al., 2021). QTL mapping summary statistics, gene lists, SNP lists, and enrichment results are provided in Supplementary Tables S1–S6.

## Supplementary Information

**Supplementary Figure S1.** Founder haplotype effects on cellular rhythm phenotypes at the mapped QTL. Phenotype values for individual mice grouped by the founder haplotype carried at the QTL marker for (**A**) rhythm amplitude at the chromosome 12 locus and (**B**) circadian phase at the chromosome 1 locus. Founder haplotypes are denoted A–H: A = A/J, B = C57BL/6J, C = 129S1/SvImJ, D = NOD/ShiLtJ, E = NZO/HlLtJ, F = CAST/EiJ, G = PWK/PhJ, and H = WSB/EiJ. Each point is one mouse; the crosshair marks the group mean ± standard deviation.

**Table S1.** Protein-coding and functional genes within the amplitude (chromosome 12; 86 genes) and phase (chromosome 1; 29 genes) QTL support intervals.

**Table S2.** Genes within suggestive QTL intervals for period (chromosomes 3, 5, X) and damping rate (chromosomes 7, 15, 16).

**Table S3.** Summary of QTL peaks: LOD scores, genomic positions, and founder allele effects for all mapped circadian phenotypes.

**Table S4.** SNP list for the amplitude QTL on chromosome 12, with founder strain genotypes and variant consequences.

**Table S5.** SNP list for the phase QTL on chromosome 1, with founder strain genotypes and variant consequences.

**Table S6.** Metascape Gene Ontology / pathway enrichment results for the 115 genes under the amplitude and phase QTL.

