## Supplementary figures and images for "Identification of Genomic Loci Associated with Cellular Rhythms in Diversity Outbred Mice"

### Supplementary Information

#### (A) Amplitude

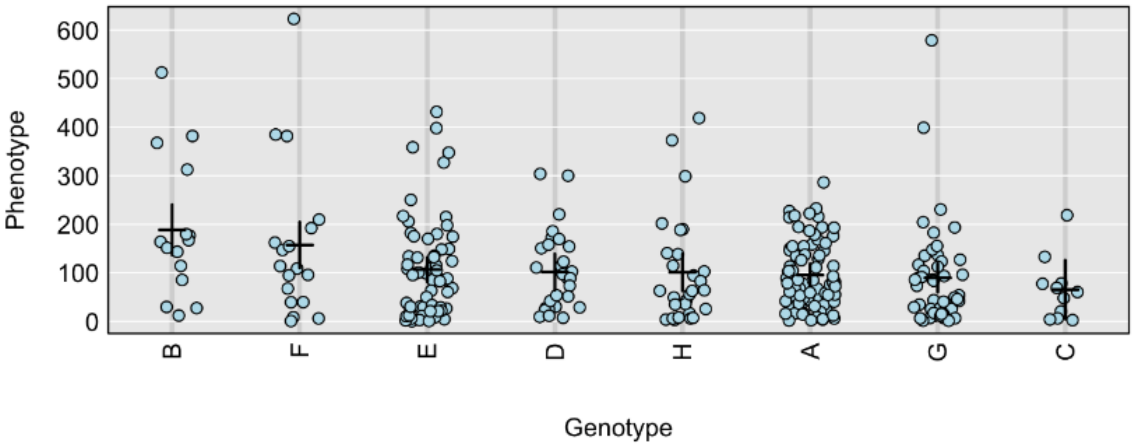


#### (B) Phase

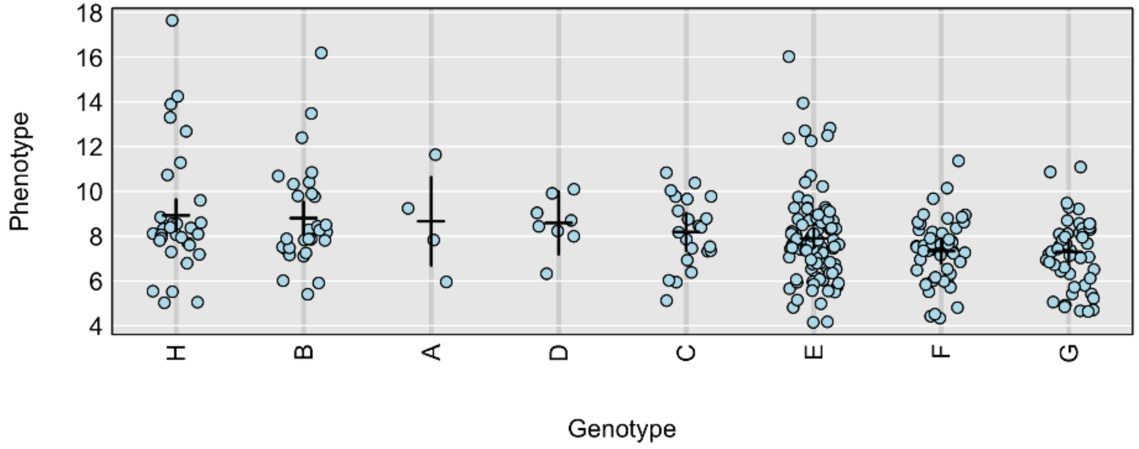
